# Substantial variation in species ages among vertebrate clades

**DOI:** 10.1101/2023.06.08.544238

**Authors:** Marcio R. Pie, Fernanda S. Caron

## Abstract

Ecological and evolutionary studies traditionally assume that species are comparable units of biodiversity. However, not only this assumption is rarely tested, but also there have been few attempts even to assess variation in most emergent, species-level traits and their corresponding underlying mechanisms. One such trait is species age, here defined as the time since the most recent common ancestor between a given species and its sister lineage. In this study, we demonstrate that different terrestrial vertebrate clades vary considerably in the age of their constituent species. In particular, species ages were youngest in mammals and birds as opposed to squamates and amphibians, although considerable variation was found within those clades as well. Sensitivity analyses showed that these results are unaffected by phylogenetic uncertainty or incomplete taxonomic sampling. Interestingly, there was little geographical correspondence in mean species age across taxa, as well as with temperature and precipitation stability over the past 21,000 years. We discuss candidate mechanisms that might explain differences in species ages among clades, and explore the implications of these findings in relation to recent advances in age-dependent speciation and extinction models of diversification.

## Introduction

Understanding the causes and consequences of variation in species numbers is at the heart of a variety of scientific disciplines, from ecology and biogeography to macroevolution. Particularly with the advent of the Linnaean classification, it is often tacitly assumed that what we mean by the word “species” is ultimately comparable across different organisms. That does not mean that interspecific differences are disregarded (e.g., [1,2]). On the contrary, understanding the causes and consequences of interspecific variability has been a major focus of ecological and evolutionary research. However, these studies tend to focus on individual traits (e.g., body size, foraging strategy, metabolic rate), rather than emergent traits at the level of the entire species. One such emergent trait is the duration of a species, given that its timescale is considerably longer than the lifetime of any particular organism.

In his classic paper, Van Valen [3] proposed the “law of constant extinction”, which states that long and short-lived taxa have equal chances of going extinct. Although some taxa indeed show age independent extinction rates (e.g., [3,4]), later papers have increasingly found departures from this rule, but the direction of such nonindependence is not congruent between taxa, with some studies showing either positive [5] or negative age-dependence [6–8] (see [9] for a review). Likewise, some authors have argued for age-dependent speciation [10]. For instance, Hagen et al. [8] showed that a model in which the rate of speciation decreases with species age was able to provide levels of clade imbalance that reflect more closely empirical trees. Although rigorous statistical approaches are increasingly designed to document age-dependent evolutionary dynamics (e.g., [11]), our understanding of general differences among clades in species ages is sorely limited. For instance, if environmental factors such as temperatureor precipitation are important drivers of variation in species ages, one would expect congruent geographical patterns in mean species ages across taxa (e.g., younger species ages in less productive regions). Alternatively, areas of particularly high climatic stability following the last glacial maximum could have led to species sorting according to their age, leading to trends in present day geographical distributions of mean species ages. To the best of our knowledge, none of these hypotheses has been explicitly tested before.

In this study, we explore several large-scale phylogenetic datasets of terrestrial vertebrates to explore variation among clades in species age, here defined as the time since the most recent common ancestor between a species and its sister lineage. In particular, species ages were youngest in mammal and bird clades as opposed to squamate and amphibian clades, although considerable variation was found between them as well.

## Materials and methods

Phylogenetic data was obtained for amphibians [12], birds [13] (Ericson backbone trees), mammals [14] (birth-death node-dated trees), and squamates [15] from VertLife.org (http://vertlife.org/phylosubsets/). The combined dataset included 32,897 species distributed across mammals (N=5,911), squamates (N=9,755), amphibians (N=7,238), and birds (N=9,993). We also split the corresponding trees into subclades to facilitate the interpretation of the results, given that the species within them tend to share similar ecologies and life-histories. The split dataset comprised 27,182 species, including mammals (Carnivora [N=334], Cetartiodactyla [N=384], Chiroptera [N=1,323], Diprotodontia [N=182], Primates [N=494)], squamates (Anguimorpha [N=257], Gekkota [N=1,638], Iguania [N=1,795], Lacertoidea [N=928], Scincoidea [N=1,757], Serpentes [N=3,560]), amphibians (Anura [N=6,416], Caudata [N=695], Gymnophiona [N=235]), and birds (Columbiformes [N=342], Passeriformes [N=6,002], Piciformes [N=450], Psittaciformes [N=390]).

Species ages were measured as the time since the most recent common ancestor between a species and its sister lineage. To account for phylogenetic uncertainty, we repeated the analyses for each of 1000 alternative topologies. We recognize that this is an imprecise measure of species’ age, given that one cannot know beforehand how long each species would last. However, it provides an operational measure of species age that is directly comparable across different taxa, even in the absence of a detailed fossil record. Also, it is possible that some clades show variation in the extent to which their species have been discovered and described, so that differences in taxonomic completeness could potentially bias estimates of species ages. To measure this potential bias, we randomly pruned each tree by a fraction of its species (1.25, 2.5, and 5%) and assess the extent to which our conclusions would change due to taxonomic incompleteness. Trees were manipulated using ‘ape’ 5.5 [16], variation in species ages across clades were visualized using ‘vioplot’ 0.3.7 [17], and significant differences were interpreted whenever groups did not overlap their confidence intervals.

Finally, we conducted a phylogenetic generalized least-squares (PGLS) regression to assess whether climate stability is a predictor of species longevity. We examined the relationship between species ages and the stability of temperature and precipitation. Paleodata on annual mean temperature and mean precipitation were obtained from PaleoView [18] and PALEO-PGEM-Series [19], covering the last 21,000 years before present (bp). To calculate statistics of climate stability over time, we utilized the ‘climateStability’ 0.1.4 package [20], which computes stability as the inverse of the mean standard deviation between time slices over the elapsed time. PGLS analyses were performed with species ages as the response variable and temperature stability, precipitation stability, and their interaction term as predictors. The analysis was repeated for a posterior distribution of 100 topologies for each taxon, considering the uncertainties in the phylogenetic relationships. In addition, we excluded species that had no spatial or climatic data (i.e., mammals [N=1,623], squamates [N=3,612], amphibians [N=1,767], and birds [N=4,009]). Geographical mapping was conducted to visualize the variables on maps, respecting the resolution of PaleoView (2.5 degrees) and PALEO-PGEM-Series (1 degree). PGLS were performed using ‘caper’ 1.0.1 [21], and maps were manipulated using ‘raster’ 3.6.20 [22], ‘rgdal’ 1.6.5 [23], and ‘sf’ 1.0.12 [24]. All analyses were carried out using R 4.1.1 [25] and QGIS [26].

## Results

There was a more than 3-fold difference in average species ages across the studied terrestrial vertebrate classes. Mammals had the youngest species (3.06 My ± 4.28, mean ± SD), whereas the oldest species were found in amphibians (10.26 My ± 11.22, mean ± SD). Sensitivity analyses performed to assess the effect of different levels of tip pruning on the inferred species ages showed minimal effects (Fig 1), indicating that the differences among groups cannot be explained by variation in taxonomic incompleteness. Interestingly, the observed differences between classes cannot simply be explained by ecto-/endothermy, given that the difference in species ages between birds and squamates on a logarithmic scale is smaller than the difference between these classes and mammals and amphibians, respectively.

**Fig 1.**
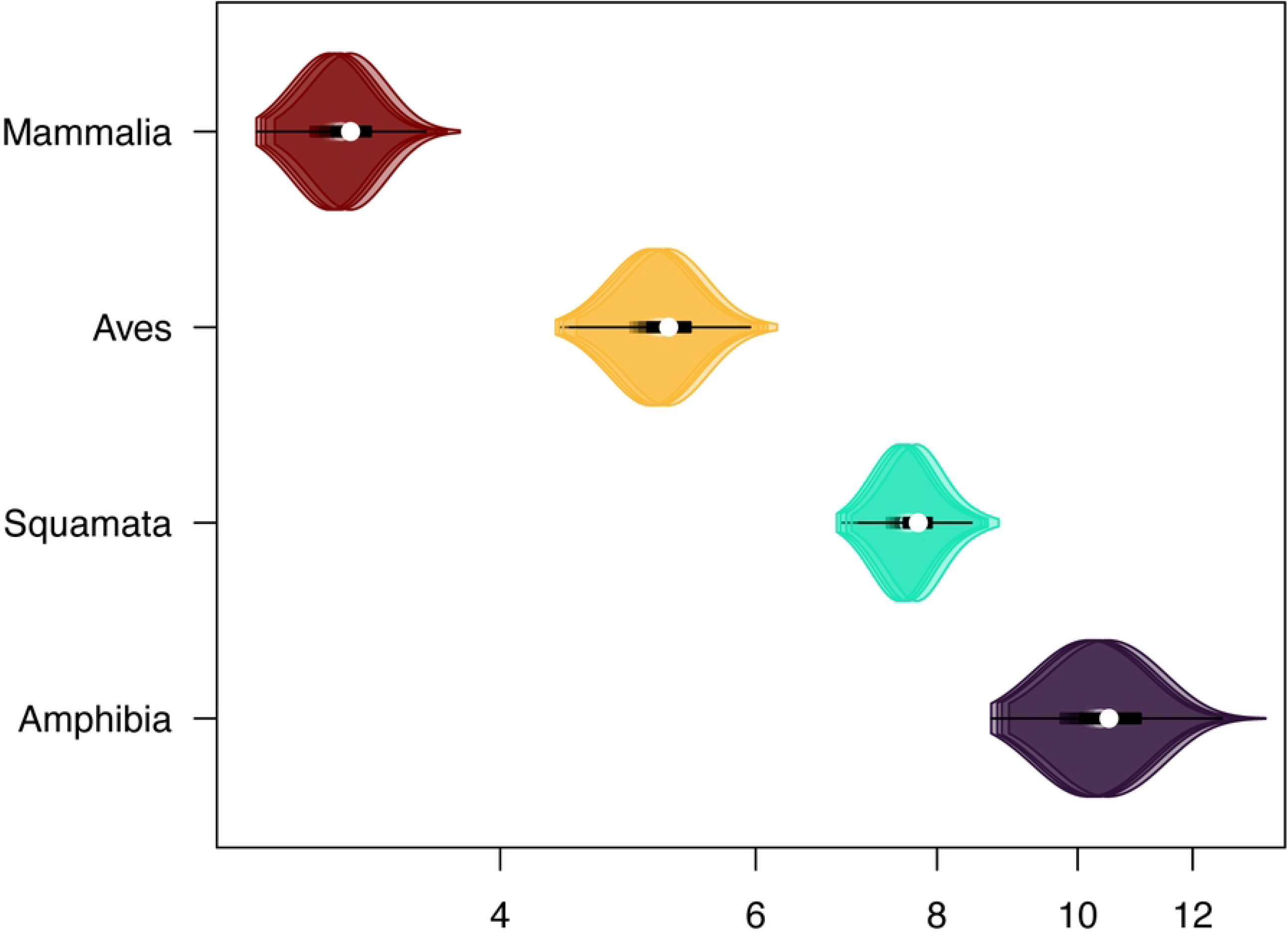
Variation in species ages across terrestrial vertebrate classes. White circles represent the median of each distribution, the thicker black lines indicate the interquartile range (IQR: Q1 and Q3 are the 25th-75th percentiles, respectively), the finer black lines span Q1-1.5 * IQR and Q3+1.5 IQR), and the width of the violin plot corresponds to the relative frequency of different values. Plots including 0, 1.25, 2.5, and 5% of randomly pruned species are overlapped to one another for each class, indicating that incomplete taxonomic sampling is unlikely to substantially affect the inferred distribution of species ages.

Interesting patterns emerged when data were divided among different subclades (Fig 2). Birds showed both the most variation within each order, but also the least variation among orders. In the case of mammals, most orders showed relatively similar species ages, except for Primates, which harbored the youngest species across all terrestrial vertebrate clades. Squamates showed relatively little variation among orders, except for snakes, which tended to display species ages comparable to birds. Finally, caecilians not only showed the oldest species ages across all terrestrial vertebrates but were also 9.84 and 10.88 My older on average than anurans and salamanders, respectively (Fig 2).

**Fig 2.**
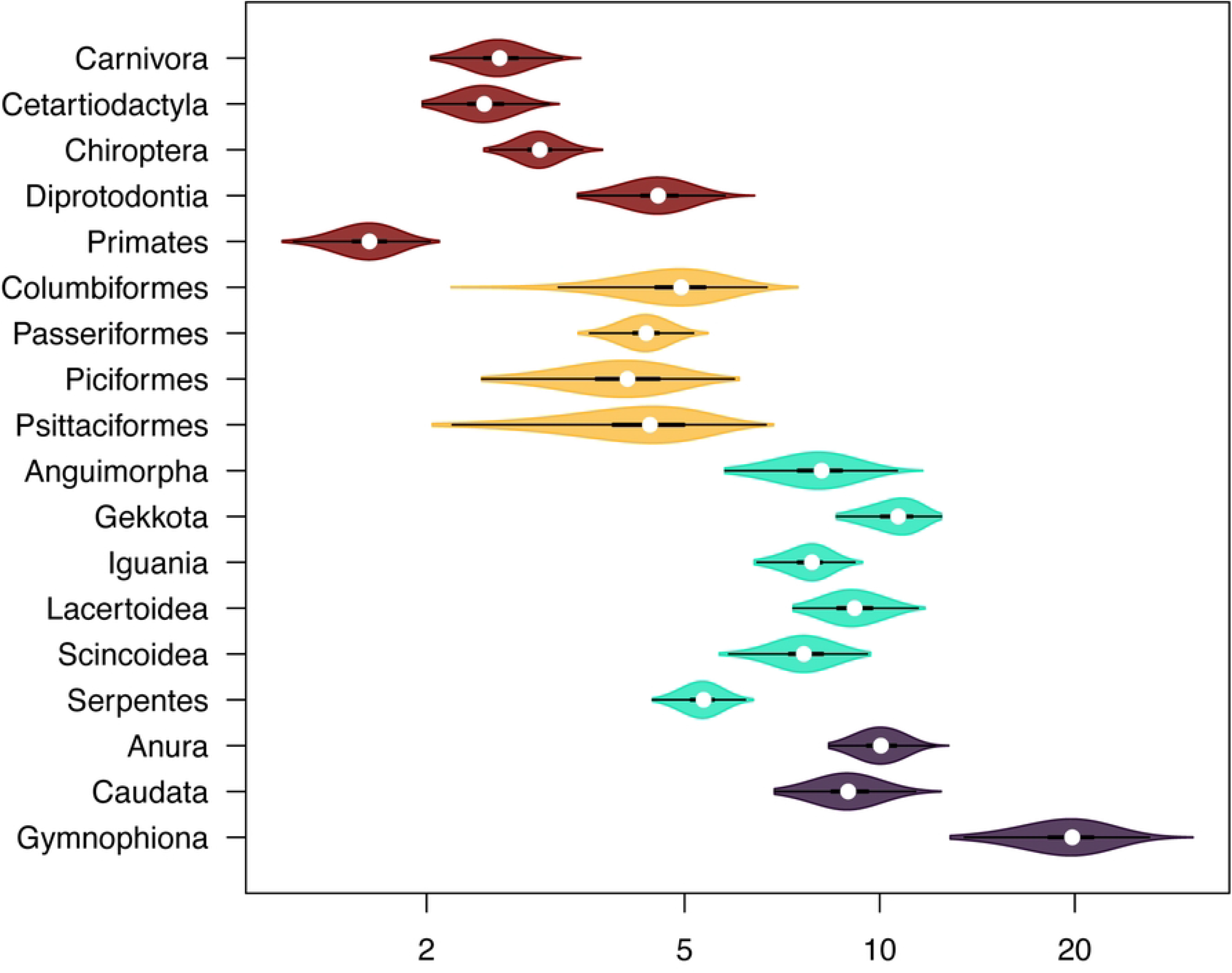
Variation in species ages across terrestrial vertebrate subclades. Each plot corresponds to the distribution across all species and alternative topologies. See legend of Fig 1 for details on the violin plots.

Geographical mapping of temperature and precipitation stability, as well as species ages, are shown in Fig 3. In both paleoclimate datasets, no clear trend can be observed between the ages of terrestrial vertebrate species and climate stability (Figs 3 and S1). These findings are further supported by the results of the PGLS analysis, where neither the taxon nor the predictors showed significance in predicting species ages (Tables 1 and S1).

**Table 1.**
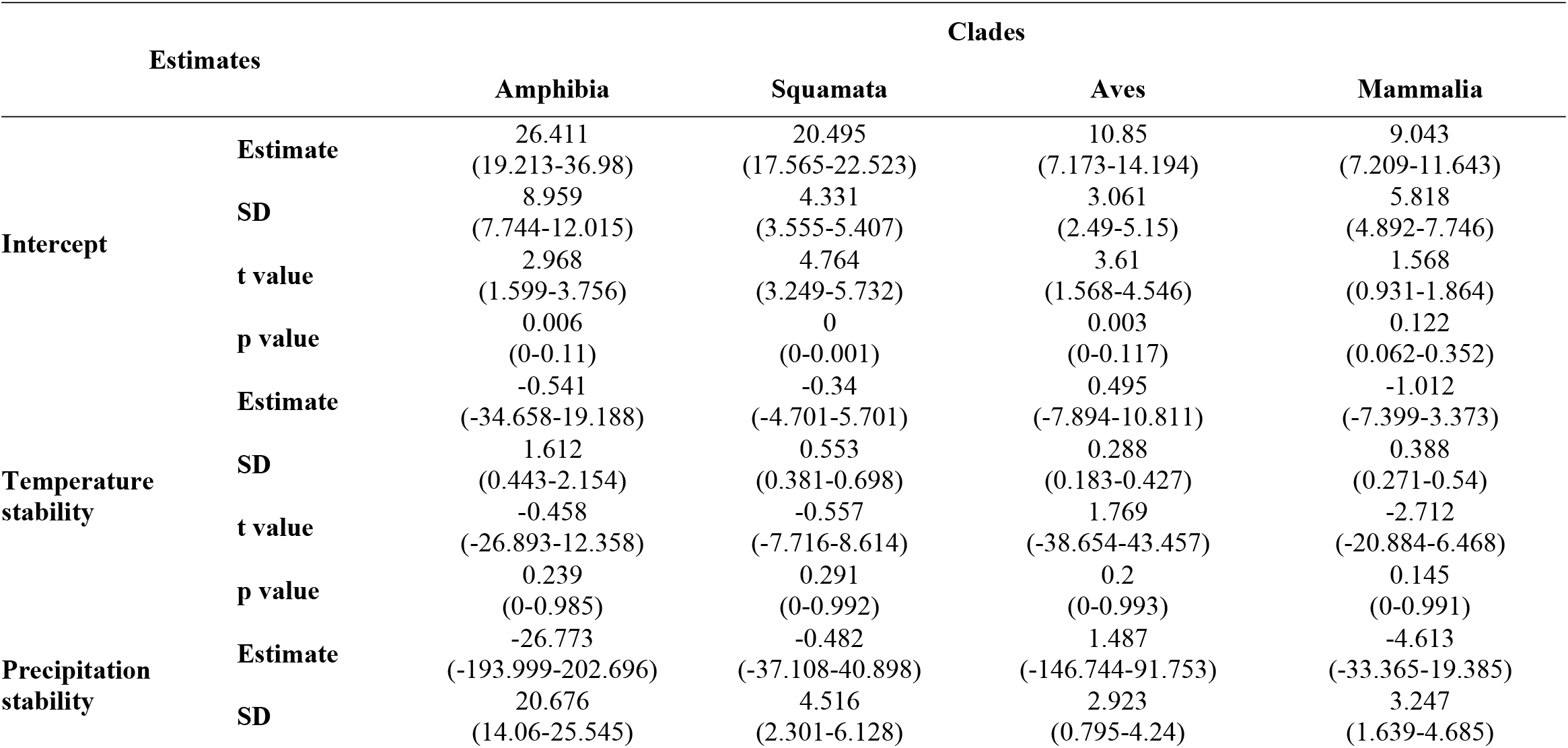
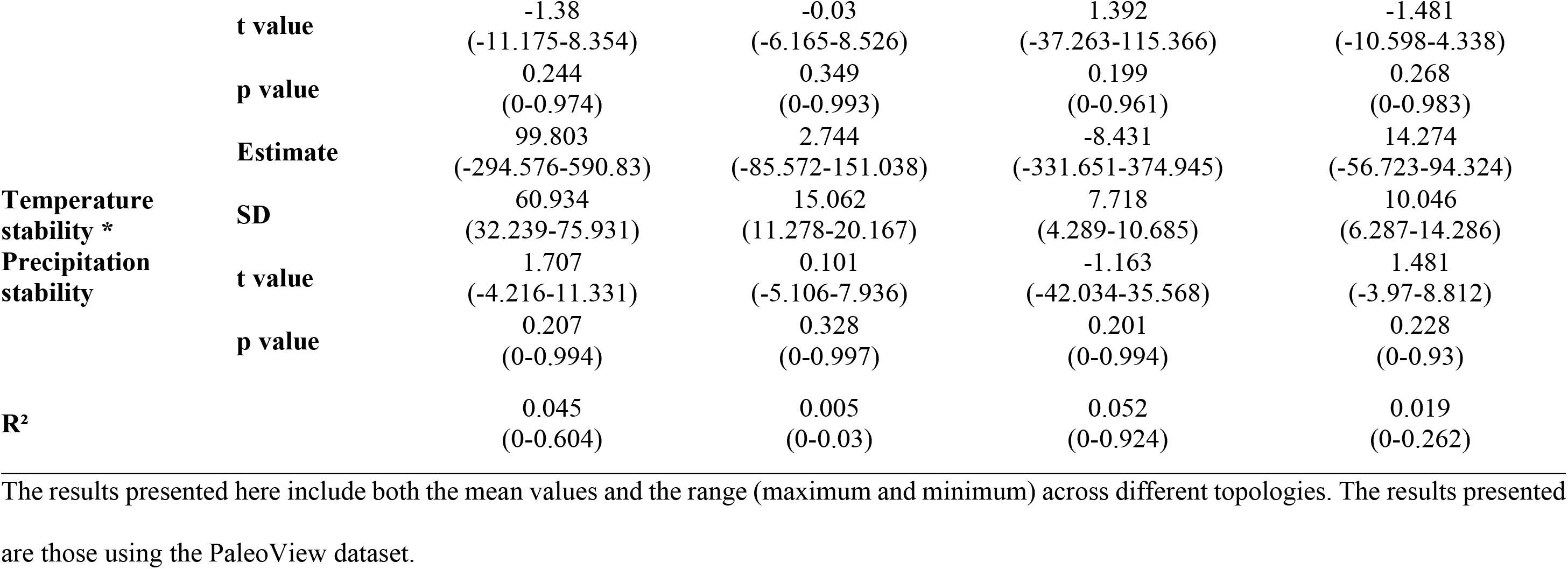
PGLS analysis performed with species ages as the response variable and temperature stability, precipitation stability, and their interaction term as predictors.

**Fig 3.**
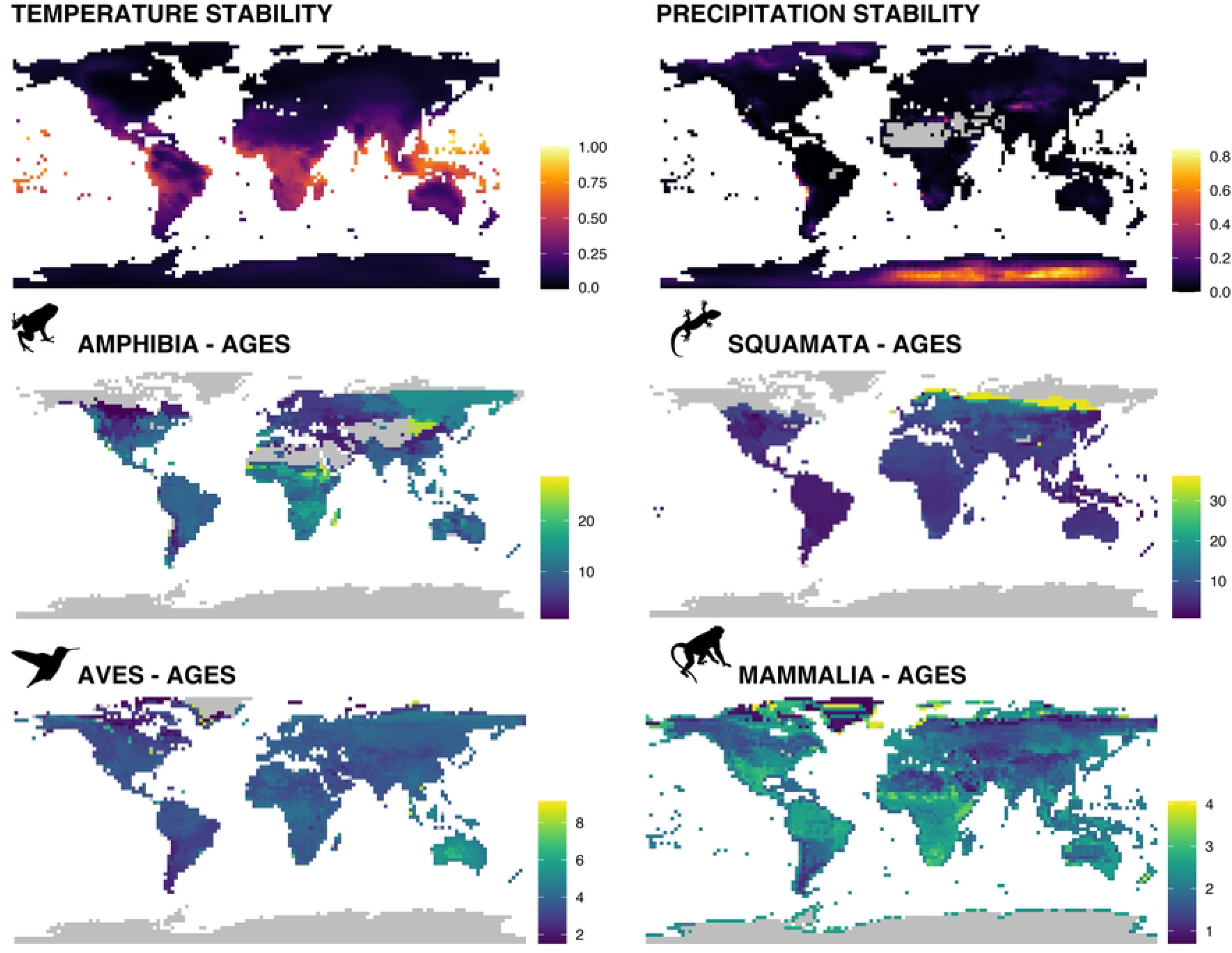
Geographical mapping of temperature and precipitation stabilities, and variation in species ages across terrestrial vertebrates. Temperature and precipitation stabilities were calculated using PaleoView dataset.

## Discussion

In this study, we uncovered substantial differences in species ages across terrestrial vertebrate clades. One must resist the temptation of disregarding these results as trivial—these organisms vary in many of their properties, why would they not vary in the age of their species as well? It is important to note that, to the best of our knowledge, there is no current framework in the literature that would predict such substantial variation based on biological first principles. For instance, one potential explanation could be variation in the rate of evolution of hybrid inviability, possibly as the result of differential rates of regulatory evolution that create developmental incompatibility. This mechanism has been argued by Fitzpatrick [27] to explain why mammals seemed to evolve complete hybrid inviability faster than birds. Although our results, in general, seem to agree that mammals indeed have relatively lower species ages than birds, there is more variation within mammals (e.g., Diprotodontia X Primates) than between mammals and birds (Fig 2), suggesting that this mechanism is unlikely to be general enough to explain our data.

Population genetic models of the speciation process have identified factors that might influence the waiting time for speciation. Some models involving stochastic peak shifts (e.g., one-locus two-allele model with underdominance [28], additive quantitative models [29]) suggest that this waiting time grows exponentially with the product of population size and the corresponding selection coefficient, leading to expected timescales that seem unrealistically long. In other words, a single peak shift resulting in strong reproductive isolation is very unlikely because the waiting time to a stochastic transition between the adaptive peaks is extremely long unless the population size is small and the adaptive valley is shallow [30]. Alternatively, the Bateson-Dobzhansky-Muller (BDM) model does not involve crossing adaptive valleys but rather only following a ridge of high fitness values [30]. In this case, the waiting time is still long (approximately the reciprocal of the mutation rate) while being independent of population size [31]. Interestingly, the addition of local adaptation to a BDM model might dramatically accelerate speciation [32], whereas adding migration might have the opposite effect [30]. We find no a priori expectation for generation time or the effect of local adaptation to be substantially different among the clades included in the present study to explain the substantial differences between them in species ages. Additionally, one might suspect that the lower dispersal capacity of many ectotherm species might lead to higher population subdivision, which in turn could accelerate speciation. However, population genetic models based on Dobzhansky-Muller incompatibilities suggest that population subdivision by itself does not affect time to speciation [33]. Interestingly, when divergence is driven by natural selection, speciation is actually faster when a species is split into two large populations [33]. On the other hand, the abovementioned models provide two non-mutually explanations. First, some of the clades might show higher dispersal rates, leading to higher migration rates which would in turn delay speciation. Second, there are substantial differences among clades in population density [34] and range size [35], which in turn could affect the probability of vicariance, as well as opportunities for local adaptation.

It is interesting to note that variation in species ages is the observation that they tend to mirror the maximum intraspecific genetic divergence. For instance, endotherms tend to show lower maximum intraspecific genetic divergence than endotherms [36]. Similar differences were found according to latitude [37] (see also [38]), which might have been influenced by the extent of past climatic variation [39]. Although there is a potential link between species ages and maximum intraspecific genetic divergence, it is important to note that those studies tended to focus only on one locus (mtDNA). Given that different loci might have distinct coalescence times, divergence at a single locus might not be representative of the entire genome. In addition, the timescales associated with locus coalescence time tend to be considerably younger than that of species ages, suggesting that a link between these two phenomena, although possible, is not necessarily obvious.

There are a few important caveats regarding our results. First, one could argue that there may be systematic variation among clades in taxonomic practice, as has been recently suggested as a factor in the recognition of the latitudinal diversity gradient [40]. Although variation across taxonomists in their willingness to describe species is indeed possible, this explanation seems unlikely, particularly at the scale necessary to explain our results. Indeed, it would mean that species limits are so ambiguous that any real biological differences would be completely swamped by taxonomic practice, and it pushes the question one step back: why would phenotypic differences that we associate with species diagnostic characters arise at different rates? Also, it is important to keep in mind that we only consider speciation by cladogenesis and not through anagenesis [41]. Although their underlying mechanisms might be different, practical limitations might mean that distinguishing between these two modes of speciation might be difficult.

In conclusion, our results uncovered an intriguing yet largely overlooked pattern across terrestrial vertebrates. Documenting such variation in other taxa and biomes, as well as assessing how the candidate mechanisms proposed here might drive interspecific variation in species ages might be a particularly exciting area for future research.

## Supporting information

**S1 Fig. Geographical mapping of temperature and precipitation stabilities, and variation in species ages across terrestrial vertebrates**. Temperature and precipitation stabilities were calculated using PALEO-PGEM-Series dataset.

**S1 Table. PGLS analysis performed with species ages as the response variable and temperature stability, precipitation stability, and their interaction term as predictors**. The results presented here include both the mean values and the range (maximum and minimum) across different topologies. The results presented are those using the PALEO-PGEM-Series dataset.

## References

1. Gutiérrez D, Wilson RJ. Intra-and interspecific variation in the responses of insect phenology to climate. J Anim Ecol. 2021;90: 248–259. doi:10.1111/1365-2656.13348

2. Olson VA, Owens IPF. Interspecific variation in the use of carotenoid-based coloration in birds: diet, life history and phylogeny: Avian carotenoid pigmentation. J Evol Biol. 2005;18: 1534–1546. doi:10.1111/j.1420-9101.2005.00940.x

3. Van Valen L. A new evolutionary law. Evol Theory. 1973;1: 1–30.

4. Pearson PN. Investigating age-dependency of species extinction rates using dynamic survivorship analysis. Hist Biol. 1995;10: 119–136. doi:10.1080/10292389509380516

5. Smits PD. Expected time-invariant effects of biological traits on mammal species duration. Proc Natl Acad Sci. 2015;112: 13015–13020. doi:10.1073/pnas.1510482112

6. Boyajian G, Lutz T. Evolution of biological complexity and its relation to taxonomic longevity in the Ammonoidea. Geology. 1992;20: 983. doi:10.1130/0091-7613(1992)020<0983:EOBCAI>2.3.CO;2

7. Condamine FL, Romieu J, Guinot G. Climate cooling and clade competition likely drove the decline of lamniform sharks. Proc Natl Acad Sci. 2019;116: 20584–20590. doi:10.1073/pnas.1902693116

8. Hagen O, Hartmann K, Steel M, Stadler T. Age-Dependent Speciation Can Explain the Shape of Empirical Phylogenies. Syst Biol. 2015;64: 432–440. doi:10.1093/sysbio/syv001

9. Januario M, Quental TB. Re-evaluation of the “law of constant extinction” for ruminants at different taxonomical scales. Evolution. 2021;75: 656–671. doi:10.1111/evo.14177

10. Liow LH, Ergon T. Fitting Ancestral Age-Dependent Speciation Models to Fossil Data. In: Allmon WD, Yacobucci MM, editors. Species and Speciation in the Fossil Record. Chicago, IL: University of Chicago Press; 2016. pp. 198–216.

11. Alexander HK, Lambert A, Stadler T. Quantifying Age-dependent Extinction from Species Phylogenies. Syst Biol. 2016;65: 35–50. doi:10.1093/sysbio/syv065

12. Jetz W, Pyron RA. The interplay of past diversification and evolutionary isolation with present imperilment across the amphibian tree of life. Nat Ecol Evol. 2018;2: 850–858. doi:10.1038/s41559-018-0515-5

13. Jetz W, Thomas GH, Joy JB, Hartmann K, Mooers AO. The global diversity of birds in space and time. Nature. 2012;491: 444–448. doi:10.1038/nature11631

14. Upham NS, Esselstyn JA, Jetz W. Inferring the mammal tree: Species-level sets of phylogenies for questions in ecology, evolution, and conservation. Tanentzap AJ, editor. PLOS Biol. 2019;17: e3000494. doi:10.1371/journal.pbio.3000494

15. Tonini JFR, Beard KH, Ferreira RB, Jetz W, Pyron RA. Fully-sampled phylogenies of squamates reveal evolutionary patterns in threat status. Biol Conserv. 2016;204: 23–31. doi:10.1016/j.biocon.2016.03.039

16. Paradis E, Schliep K. ape 5.0: an environment for modern phylogenetics and evolutionary analyses in R. Schwartz R, editor. Bioinformatics. 2019;35: 526–528. doi:10.1093/bioinformatics/bty633

17. Adler D, Kelly ST. vioplot: violin plot. R package version 0.3.7. 2021. Available: https://github.com/TomKellyGenetics/vioplot.

18. Fordham DA, Saltré F, Haythorne S, Wigley TML, Otto-Bliesner BL, Chan KC, et al. PaleoView: a tool for generating continuous climate projections spanning the last 21 000 years at regional and global scales. Ecography. 2017;40: 1348–1358. doi:10.1111/ecog.03031

19. Barreto E, Holden PB, Edwards NR, Rangel TF. PALEO-PGEM-Series: A spatial time series of the global climate over the last 5 million years (Plio-Pleistocene). Glob Ecol Biogeogr. 2023; geb.13683. doi:10.1111/geb.13683

20. Owens HL, Guralnick R. climateStability: An R package to estimate climate stability from time-slice climatologies. Biodivers Inform. 2019;14: 8–13. doi:10.17161/bi.v14i0.9786

21. Orme D, Freckleton R, Thomas G, Petzoldt T, Fritz S, Isaac N, et al. caper: comparative analyses of phylogenetics and evolution in R. R package version 1.0.1. 2018. Available: https://CRAN.R-project.org/package=caper.

22. Hijmans RJ, van Etten J, Sumner M, Cheng J, Baston D, Bevan A, et al. raster: Geographic Data Analysis and Modeling. R package version 3.6-20. 2023. Available: http://CRAN.R-project.org/package=raster

23. Bivand R, Keitt T, Rowlingson B, Pebesma E, Sumner M, Hijmans R, et al. rgdal: Bindings for the “Geospatial” Data Abstraction Library. R package version 1.6-5. 2023. Available: http://cran.r-project.org/package=rgdal

24. Pebesma E. Simple Features for R: Standardized Support for Spatial Vector Data. R J. 2018;10: 439. doi:10.32614/RJ-2018-009

25. R Core Team. R: A language and environment for statistical computing. Version 4.1.1. 2021. Available: https://www.R-project.org/.

26. QGIS Team. QGIS Geographic Information System. Open Source Geospatial Foundation Project. 2023. Available: http://qgis.osgeo.org

27. Fitzpatrick BM. Rates of evolution of hybrid inviability in birds and mammals. Evolution. 2004;58: 1865–1870. doi:10.1111/j.0014-3820.2004.tb00471.x

28. Lande R. Effective deme sizes during long-term evolution estimated from rates of chromosomal rearrangement. Evolution. 1979;33: 234–251. doi:10.1111/j.1558-5646.1979.tb04678.x

29. Barton NH, Charlesworth B. Genetic Revolutions, Founder Effects, and Speciation. Annu Rev Ecol Syst. 1984;15: 133–164. doi:10.1146/annurev.es.15.110184.001025

30. Gavrilets S. Pterspective: Models of speciation: What have we learned in 40 years?Evolution. 2003;57: 2197–2215. doi:10.1111/j.0014-3820.2003.tb00233.x

31. Nei M. Mathematical models of speciation and genetic distance. In: Karlin S, Nevo E, editors. Population Genetics & Ecology. New York, NY: Academic Press; 1976. pp. 723–768.

32. Schluter D. The ecology of adaptive radiation. Oxford: Oxford Univ Press; 2000.

33. Orr HA, Orr LH. Waiting for speciation: The effect of population subdivision on the time to speciation. Evolution. 1996;50: 1742–1749. doi:10.1111/j.1558-5646.1996.tb03561.x

34. Pie MR, Caron FS, Divieso R. The evolution of species abundances in terrestrial vertebrates. J Zool Syst Evol Res. 2021;59: 2562–2570. doi:10.1111/jzs.12526

35. Pie MR, Divieso R, Caron FS. Do Geographic Range Sizes Evolve Faster in Endotherms? Evol Biol. 2021;48: 286–292. doi:10.1007/s11692-021-09537-x

36. Tingley R, Dubey S. Disparity in the timing of vertebrate diversification events between the northern and southern hemispheres. BMC Evol Biol. 2012;12: 244. doi:10.1186/1471-2148-12-244

37. Cattin L, Schuerch J, Salamin N, Dubey S. Why are some species older than others? A large-scale study of vertebrates. BMC Evol Biol. 2016;16: 90. doi:10.1186/s12862-016-0646-8

38. Weir JT, Schluter D. The Latitudinal Gradient in Recent Speciation and Extinction Rates of Birds and Mammals. Science. 2007;315: 1574–1576. doi:10.1126/science.1135590

39. Dubey S, Shine R. Geographic variation in the age of temperate-zone reptile and amphibian species: Southern Hemisphere species are older. Biol Lett. 2011;7: 96–97. doi:10.1098/rsbl.2010.0557

40. Freeman BG, Pennell MW. The latitudinal taxonomy gradient. Trends Ecol Evol. 2021;36: 778–786. doi:10.1016/j.tree.2021.05.003

41. Ezard THG, Pearson PN, Aze T, Purvis A. The meaning of birth and death (in macroevolutionary birth–death models). Biol Lett. 2012;8: 139–142. doi:10.1098/rsbl.2011.0699

